## Supplementary Information for "A versatile active learning workflow for optimization of genetic and metabolic networks"

**Page**

| | |
|---|---|
| Supplementary Notes 1-7 | 1-8 |
| Supplementary Tables 1-7 | 9-12 |
| Supplementary Figures 1-20 | 13-29 |
| Supplementary References | 30 |

Supplementary Notes:

**Supplementary Note 1**: Active learning process for optimization of cell-free Gfp production
We started the active learning cycle by generating 20 random compositions for the first round, pipetting them, measuring the Gfp fluorescence in a plate reader, then we collected the results after 6 hours of incubation at 30 °C. The mean and standard deviation of each composition was imported to the model, and after training, it suggested a new set of 20 compositions. The yield was calculated from the measured Gfp fluorescence of every composition normalized by a composition in which all variable factors are at their mid-range. This normalization was used for active learning (**Data availability**). In the end, as plotted in **Fig. 1e**, all values were normalized by composition from a common protocol in the field of cell-free synthetic biology to be comparable with other studies (**Methods**). The cycle (**Fig. 1d**) was repeated for 10 rounds. In each round, the model became more predictive in generating new compositions and ranking the best 20 suggestions for the next round. Since pipetting 13 elements in 20 samples is error-prone and needs a substantial amount of time and effort, we developed a table-to-speech virtual assistant that is run on Google Colab and can be easily used on a computer or smartphone. This tool takes as input the suggested table of volumes to pipette, goes through it line by line (factors), ranks them from minimum to maximum volume, and reads and graphically shows them on the screen. With organized pipette tip sets and destination tubes (**Methods**), this considerably improved the speed, accuracy, and comfort of pipetting of such complex compositions.

**Supplementary Note 2**: User guide and description of features of the modular workflow
**Fig. 2b** shows the workflow of using METIS from the adjustment of the parameters to data visualization and analysis. The user input consists of two sections for i) active learning parameters and ii) factors with a range and/or category. In the active learning parameters section, the user should define the number of combinations willing to perform in each round depending on the number of factors and their conditions, also considering the equipment for i) pipetting compositions (or cloning genetic constructs in case of biological sequences) and ii) measuring the objective function. The number of rounds of active learning also should be defined as any arbitrary number. It is important to note that the number of total experiments is recommended to be divided into more rounds as this improves the power of active learning[1]. Other parameters are the total volume of each composition, the portion of the volume that could be varied excluding and listing the constant factors. This is important to specify the total volume and the volume of fixed components subtracted from it because the model needs to convert concentrations to volumes and vice versa. The minimum droplet size defines the minimum volume unit of liquid handling robots or the minimum unit that one can pipette, usually 0.2 or 0.5 µL. This means all the volumes proposed by the model will be a factor of 0.2/0.5 µL or, for example, 2.5/25 nL with Echo® acoustic robot.

The exploration/exploitation ratio referred to as “exploration” in the parameters section is one of the most important elements to thoughtfully define, especially when the user applies the workflow for a different application than those presented in this work. For applications with numerical factors (i.e., our examples in **Fig. 1, 3**), the exploration rate should be >1 for the early rounds of active learning, and toward the end of the cycle, it should tend to values <1. Importantly, these values depend on the number, importance, and type (numerical or categorical) of factors, therefore, for tailoring the workflow, it should be taken into account. A high exploration indicates that the model will take more risks in suggesting combinations (exploration) rather than being efficient (exploitation). High exploration ratios are needed in the early rounds to explore the space and escape from local optimal combinations. However, in the later rounds, more efficient suggestions are required, which corresponds to focusing on achieving higher yields using what the model has learned. The ratio should be very carefully assigned, and the users should rely on their knowledge of the system or check a few of the suggestions, especially for the use with categorical factors like for our examples in **Fig. 4, Supplementary Fig. 19**. Note that, even if an exploration ratio is assigned for Day 1 (in the absence of Day 0), the first step is fully randomized, and the ratio has no effect on it. The exploration ratio for the examples reported in this work is as follows. 10 rounds of Gfp production in the cell-free system (**Fig. 1e**): Not defined, 1.41, 1, 1, 1, 1, 0.5, 0.5, 0.5, for 10 rounds of *LacI* gene circuit optimization (**Fig. 3c**): Not defined, 1.41, 1.0, 1.0, 1.41, 1.41, 1.41, 1.41, 1.0, 0.5, for 2 rounds of 20 most informative combinations of *LacI* gene circuits with purified LacI (**Fig. 3i**): Not defined for Day 0, 0.5 for Day 1, for 4 rounds for the transcription and translation unit (**Fig. 4c**): Not defined, 1.41, 4.0, 1.0, for 2 rounds of 20 most informative combinations of transcription translation unit *in vivo* (**Fig. 4f**): Not defined for Day 0, 2 for Day 1, and for the single round of enzyme engineering suggestions for the hypothetical next day (**Supplementary Fig. 20**): 2.0. It must be noted that, since in METIS optimization combinations are sorted based on only their std value and exploitation is set to zero (this enables picking the most uncertain combination for the next round (**Methods**)), the exploration ratio is not defined.

In the factors section (**Fig. 2b**), the user should list all the elements participating in the objective function, that can be categorical and/or numerical features. Numerical features are defined by values, such as the concentration of factors in an *in vitro* system or in a growth medium, or strength of regulatory sequences such as promoters, RBSs (ribosome binding site), or CRISPR RNAs. Categorical features are not defined by values but characters/names such as regulatory sequences without numerical scores, or when for a gene there are multiple candidates from different organisms. Combined features are those with categories (alternative sequences/genes/constructs) and their level through concentration or strength (*LacI* gene circuits in **Fig. 3**). The user can range numerical values either by giving the range boundaries (minimum and maximum) or by specifying all the values. After running this section, for factors with range boundaries, it might be the case that the number of conditions is high, or their distribution lacks the desired coverage within the range. In this situation, we suggest picking 5 to 10 values from the proposed conditions from the output code of this section and manually specifying those factors' concentrations. The stocks' concentration is an important parameter since it must support the volume of the minimum final concentration in the defined range dependent on the minimum volume unit. One highlighted feature of our workflow that empowers its performance is letting the range of factors have more variable values than only 4 conditions in Borkowski *et al*[2]. The more randomized space allows the model to sample more informatively within the ranges, and this flexibility improves the performance. After the user has defined the active learning parameters and factors, the model shows all ranges taking into account the final volume of the mixture to give a percentage of possible compositions. A percentage of 100 means it is possible to compose a condition in which all factors are at their highest volume (concentration) and lower percentages indicate compositions are limited. When needed, the stocks' concentration can be changed, and if on the other hand it limits the lowest volume, two stocks with different concentrations can be provided. This is one of the features of the workflow that can solve issues when it comes to practice.

If a pre-existing dataset is used to train the model prior to starting active learning, they can be imported (compositions and yields) to initiate the workflow on Day 0 instead of Day 1 (used examples for 20 most informative combinations in **Fig. 3i, 4f** and the enzyme engineering application in **Supplementary Note 7, Supplementary Fig. 19, 20**). The dataset should be provided in the format of the results.csv files (**Data availability**).

When the parameters and factors are all set up, the model can generate the first round of experiments from the file Volumes_1.csv (round 1). The model generates Volumes.csv and

Concentration.csv files in each round. The user should perform the suggested experiments, measure the objective function, and insert the results (mean and standard deviations) into new columns in the Concentrations_1.csv file generated by the model for that round and rename it to Results_1.scv. To continue with the next day (round), the user should upload the Results_1.csv file into the Google Colab notebook, run all sections, and the model executes the next day's compositions. In this step, the model trains itself on the results and suggests new compositions predicted to give higher values for the objective function (or more predictive combinations in METIS prediction to build a more predictive model and not necessarily optimizing the objective function). This cycle is repeated for n that is defined in the parameters section. The file used for pipetting is Volume.csv generated in each round. If the laboratory accesses Echo® acoustic liquid handler, our workflow can generate an input file corresponding to the suggested compositions. In this case, the source and destination plate wells should be specified by the user after downloading the Echo® file. Otherwise, compositions can be pipetted by other liquid handlers or manually using the Volume.csv file. In examples in **Fig. 1, 3**, for 20 compositions in each round, it took us around three hours to pipette all compositions using our table-to-speech virtual assistant. The table-to-speech assistant increases the speed, accuracy, and ease of pipetting that should be used through a separate Google Colab notebook that we also provided in this work. The notebook takes the Table2Speech_Volume.csv file as input, ranks, and reads them throughout all compositions for each factor. By adjusting the parameters in Google Colab the assistant will go to the next well with any click specified by the user (**Supplementary Note 1, Code availability**). If the objective function is as in our examples in **Fig. 4, Supplementary Fig. 20**, the combinations should be constructed or cloned, the Volume.csv lists the categories of factors that should be constructed and not real volumes to pipette.

During active learning, parameters or ranges can be altered at any round. This feature enables the user to make readjustments for the next rounds depending on the model's performance, the evolution of the objective function over rounds, and how factors behave within their ranges. Additionally, in the section named "specials", customized compositions can be manually added to any round and these will be subtracted from the number of suggestions in that round. If a user is willing to readjust or import special compositions, we recommend doing this in the later rounds of active learning and let the model perform its task in early rounds. For the *LacI* circuit optimization cycle, we show examples where customized compositions have been manually added. Controls (i.e., negative control or reference combination) could be part of specials, in particular when an Echo□ file is used.

The workflow generates cumulative outputs in each round (see **Methods** for how they are mathematically/statistically generated). The first set of outputs are compositions as Concentrations.csv and Volumes.csv. The second set is the list of K most informative combinations. This list can be used after active learning, to facilitate optimization of similar systems with the same type of factors and objective function such that one does not need to redo the whole active learning. By doing experiments for K (user-defined) combinations and measuring their objective function, results are imported as Day 0 to optionally continue with one or more rounds of active learning (see *LacI* circuit section). The last set of outputs are several types of analysis/visualization and the respective raw data, a box plot for the evolution of objective function over rounds, two groups of individual plots for each factor one showing the daily variation of factors within their ranges, and the other group is about all measured yields within ranges. The other analysis/visualization outputs are a plot and list of feature importance values, contribution of each factor in the prediction of yield by the model. These provide information on factors playing more important roles and which have less or no effect on the objective function. The last module is a heatmap of mutual interactions between every two factors providing useful results to optimize and study the system as well as to spotlight unknown interactions.

**Supplementary Note 3**: Complementary note on the experiment and discussion for optimization of *LacI* gene circuits

The median of objective function increased especially from Day 6, and the fold-change raised from less than 4 to over 8 on Day 10 (**Fig. 3c**). The median on Day 9 dropped because we included 5 of 20 as special compositions with high concentrations of $P_{T7}$-*LacI* plasmid and purified T7 RNA polymerase that led to very low objective function values. We did so since the model was trending to suggest very low concentrations of these two factors (**Supplementary Fig. 8**) which we first assumed would improve the fold change and objective function. Although this manipulation in the active learning process reduced the objective function values on Day 9, we gained a more profound insight into the system's behavior. Additionally, this helped the model to improve the objective function values at the highest on Day 10. We hypothesize that the low protein production is because of resource competition or inhibitory protein-protein interactions especially at high concentrations of the plasmid that higher amounts of amino acids and tRNAs could not compensate for it. This was noticed by the model such that during the 10-day period it was tending to suggest lower concentrations of $P_{T7}$-*LacI* plasmid and T7 RNA polymerase (**Fig. 3d, Supplementary Fig. 8**). **Fig. 3d, e** represent yield data points within each factor's range and the features' importance, respectively. The model calculated that $P_{T7}$-*LacI* plasmid is by far the most important feature because of its hugely negative effect on protein production (**Fig. 3e**).

To test the hypothesis of resource competition or inhibitory protein-protein interaction, we designed titration experiments in which with the optimal composition from active learning (which was with the pTHS circuit) we varied the concentration of $P_{T7}$-*LacI* plasmid and T7 RNA polymerase (**Fig. 3f**). Increasing the concentration of $P_{T7}$-*LacI* plasmid or T7 RNA polymerase diminishes the fold-change × dynamic range (FC × DR) and fold change (FC). The maximum of these values was achieved at low concentrations of $P_{T7}$-*LacI* plasmid and the highest amount of T7 RNA polymerase. To further support this hypothesis, instead of pTHS circuit, we added a *Gfp* expressing plasmid with a constitutive promoter that has no interaction with the $P_{T7}$-*LacI* plasmid and T7 RNA polymerase but shares the same resources for transcription and translation. Increasing the concentration of either $P_{T7}$-*LacI* or T7 RNA polymerase depleted resources for expression of the constitutive *Gfp* (**Fig. 3g**). Since the addition of the second plasmid prevented reaching a sufficient production-repression balance, we sought to test the system with His-tag purified LacI. We extracted the 20 most informative combinations from the above active learning cycle. We removed T7 RNA polymerase from the factors and replaced the $P_{T7}$-*LacI* plasmid with His-tag purified LacI and gave random values to it (**Fig. 3h**). These 20 combinations were performed, the results were collected and imported as Day 0 to the workflow. We already could see a huge improvement in the objective function and fold-change on Day 0 (**Fig. 3i**) due to using purified LacI. As high yield values were associated with the lower concentrations of purified LacI, we widened the range of purified LacI toward lower concentrations for the next round (Day 1). The Day 1 experiment resulted in 4 data points with a fold change of around 100 (right plot in **Fig. 3i**).

**Supplementary Note 4**: Setup for the METIS based optimization of the CETCH cycle

For the optimization of the CETCH cycle, we did 125 conditions in triplicates per round. In each condition, all components were variable except for the substrate propionyl-CoA, which was fixed to 100 µM. For the buffer (Hepes), we not only wanted to optimize the concentration, but also test different pH. Therefore, we gave the possibility to choose between 6 different pHs (7.0, 7.2, 7.4, 7.6, 7.8 and 8.0). For the five rounds of unrestricted optimization, we used exploration values of 1.41, 1.41, 1.0, 1.0 and 0.5. After transforming the data for the efficiency optimization ($[Glycolate]/[Enz_{tot}]$= Gycolate yield in µM and total concentration of enzymes in µM), we did three additional rounds of efficiency optimization where we used exploration values of 1.0, 0.5 and 0.5.

After the first two rounds, we purified new batches of mco, hbd, cat and ssr. In the third round, using the new enzyme stocks, we noticed that our positive control (see material and methods) had a roughly three times higher product yield compared to the first two rounds. Therefore, we tested our control manually with the new enzymes and 4 control setups where only the old enzyme stock of either mco, hbd, cat or ssr was used. We could identify the old catalase stock as the reason for the lower yields in the first two rounds (**Supplementary Fig. 16b**). We stored the new catalase stock in liquid nitrogen upon use. Additionally, we removed unnecessary values after the first three rounds to reduce the combinatorial space (see **Supplementary Table 2**). All other values are in the GitHub repository.

Assays for determination of new enzyme stocks after round two (**Supplementary Fig. 16b**)
The assays were done in triplicates containing 30 µL volume each and were carried out in a 1.5 mL reaction tube (at 30 °C, 500 rpm). The reactions were started with 100 µM propionyl-CoA. 8 µL samples were taken and quenched in 1 µL 50% formic acid and 1 µL 500 mM sodium polyphosphate (emplura®) at 1, 2, and 3 h. The samples were spun for 20 min at 4 °C and 20.000g, before the supernatant was transferred into new tubes. As described earlier, the samples were diluted and mixed with the internal standard for LC-MS measurement.
The assays contained:
100 mM HEPES pH 7.6, 5 mM $MgCl_2$, 50 mM sodium bicarbonate, 20 mM sodium formate, 20 mM creatine phosphate, 0.5 mM CoA, 0.1 mM $CoB_{12}$, 2 mM ATP, 5 mM NADPH, 3.06 µM pco, 0.62 µM ccr, 0.74 µM epi, 0.30 µM mcm, 2.62 µM scr, 0.53 µM ssr, 5.34 µM hbs, 0.73 µM hbd, 0.58 µM ecm, 21.90 µM mco, 0.28 µM mch, 2.79 µM mcl, 1.64 µM cat, 14.56 fdh, 0.02 µM ca, 1.10 µM gor, 0.39 µM ck.
The experiments labeled mco, hbd, cat and ssr were done with the old stocks of the enzymes used in the first two rounds. The controls contained enzymes from the four new enzymes batches.

Assays for different concentrations of hbs and cobalt (**Supplementary Fig. 16a**)
The assays were done in triplicates containing 30 µL volume each and were carried out in a 1.5 mL reaction tube (at 30 °C, 500 rpm). The reactions were started with 100 µM propionyl-CoA. 8 µL samples were taken and quenched in 1 µL 50% formic acid and 1 µL 500 mM sodium polyphosphate (emplura®) at 1, 2 and 3 h. The samples were spun for 20 min at 4 °C and 20.000g, before the supernatant was transferred into new tubes. As described earlier, the samples were diluted and mixed with the internal standard for LC-MS measurement.

The hbs assays contained:
100 mM HEPES pH 7.6, 5 mM $MgCl_2$, 50 mM sodium bicarbonate, 20 mM creatine phosphate, 0.5 mM CoA, 0.1 mM $CoB_{12}$, 2 mM ATP, 10 mM NADPH, 10 mM NADH, 3.06 µM pco, 0.62 µM ccr, 0.74 µM epi, 0.30 µM mcm, 2.62 µM scr, 0.53 µM ssr, {0.53 µM hbs (10%), 5.34 µM hbs (control), 26.7 µM hbs (500%)}, 0.73 µM hbd, 0.58 µM ecm, 21.90 µM mco, 0.28 µM mch, 2.79 µM mcl, 1.64 µM cat, 0.02 µM ca, 1.10 µM gor, 0.39 µM ck.

The cobalt assays contained:
100 mM HEPES pH 7.6, 5 mM $MgCl_2$, 50 mM sodium bicarbonate, 20 mM sodium formate, 20 mM creatine phosphate, 0.5 mM CoA, {1.0 mM cobalt (1 mM), 0.0 mM cobalt (0 mM)}, 2 mM ATP, 5 mM NADPH, 3.06 µM pco, 0.62 µM ccr, 0.74 µM epi, 0.30 µM mcm, 2.62 µM scr, 0.53 µM ssr, 5.34 µM hbs, 0.73 µM hbd, 0.58 µM ecm, 21.90 µM mco, 0.28 µM mch, 2.79 µM mcl, 1.64 µM cat, 14.56 fdh, 0.02 µM ca, 1.10 µM gor, 0.39 µM ck.

**Supplementary Note 5**: Optimization versus prediction purpose

So far, we demonstrated the use of our active learning workflow for the experimentally guided optimization of cell-free protein expression, *LacI* gene circuit, transcription and translation unit, and the CETCH cycle. Machine learning algorithms learn patterns in data and predict unseen cases which can be either applied for optimization (as demonstrated with our workflow above) or prediction purposes. Note that while both approaches employ a dedicated prediction step, their overall aim is different (**Supplementary Table 1**). To further modularize our workflow, we created an additional METIS package for prediction (not optimization) of an objective function of given combinations. **Supplementary Table 1** summarizes the features of two Google Colab packages, METIS optimization and METIS prediction, which differ in goal and application, query strategy, and outputs. The query strategy or the approach of METIS optimization is maximizing the objective function as well as the standard deviation of the ensemble regressor (i.e., the result of multiple regressors that individually operate on the data) whereas METIS prediction only maximizes the standard deviation of the ensemble regressor. These two also differ in the output types; METIS optimization suggests combinations with high ranked objective function whereas METIS prediction generates a trained model that computes the objective function for input combinations.

Each round's suggestions in METIS prediction aim to build a more predictive model by maximizing the correlation between predicted and measured objective functions (and not necessarily maximizing the objective function as in METIS optimization), hence improve the $R^2$ of prediction over rounds of learning. Beyond these differences, the two notebooks share the same features/modules as described in **Fig. 2** and **Supplementary Note 2**. To test the prediction notebook, we ran a simulation on a dataset of 1094 data points[3] from the PURE (purified recombinant elements) cell-free protein expression system in which recombinant proteins and the buffer composition were varied (**Supplementary Note 6, Supplementary Fig. xx**). To demonstrate the performance of the model, we split the dataset into train and validation sets and used the validation set to measure (using $R^2$) the accuracy of the prediction on the test set. Because of the random nature of splitting, the simulation was repeated 5 times. (**Supplementary Fig. 18**). Overall, the workflow was able to improve the prediction of the objective function over rounds of active learning.

**Supplementary Note 6**: <u>Application of METIS prediction for simulation on a PURE cell-free system dataset</u>

To show METIS's potential in predicting an objective function, we performed an active learning simulation on an available PURE (purified recombinant elements) cell-free system dataset[3] (**Supplementary Fig. 18a, b**). We also assessed the predictability of the dataset and our model's general performance, we calculated 5-fold cross-validation on the whole dataset (1098 data points). The result of 5-fold cross-validation and the prediction of a single sample test set is shown in **Supplementary Fig. 18c, d**, respectively.

We simulated an active learning cycle as presented in **Supplementary Fig. 18a**:

- 20% of the dataset was separated as the test set to be used for validation, which was not taken into account when the model is trained on the training set (the other 80%).
- In the first round, 80 data points from the training dataset were selected randomly and the model was trained on them.
- In the next round, using what the model has learned, the rest of the training set (80 subtracted) was sorted based on the uncertainty value of combinations (**Methods**) and the top 80 combinations were picked.
- The yield of the picked combinations was assigned from the dataset and the model was trained on them.
- This process was repeated 10 times, hence, at the end the model was trained on 800 data points.

- In each step performance of the model was evaluated on the test set.

This process includes multiple random steps which is why we repeated the whole process five times as shown in **Supplementary Fig 18a**.

**Supplementary Note 7**: Application of the workflow for combinatorial enzyme engineering
Machine learning is a powerful tool to address complicated biological questions such as prediction/engineering of the structure/activity of proteins/enzymes[4,5]. Challenges in engineering a protein using machine learning tools are the dependency on large datasets and difficulties to make genotype-phenotype links for hundreds or thousands of variants[4]. Except for phenotypes easy to measure such as those related to regulatory sequences (transcription factors), there is a lack of modular characterization methods for engineering proteins. Enzymes are of difficult proteins to engineer because each enzyme requires a different characterization method which mainly allows for low to medium throughput experiments. Moreover, traditional enzyme engineering approaches which mostly rely on altering one amino acid at a time, are likely to be trapped into the local optima of the enzyme activity[6]. Since our tool is able to work with minimal datasets and the gradient boosting algorithm can capture mutual interactions, it can also be used for engineering enzymes. From a recent study in our lab, we took a dataset of mutants of oxalyl-CoA decarboxylase from *Methylorubrum extorquens*[7] to run simulations and give an example of how to use the workflow for such applications. **Supplementary Fig. 19a** shows the active site of the enzyme and surrounding amino acid residues.

We imported the dataset comprising 847 combinations of mutations in the active site of the enzyme (**Data availability**). As provided in the modular workflow, we first extracted and plotted the feature importance values (**Supplementary Fig. 19b**). E135G and Y497F were calculated as conditions with the highest effect on the yield. Although such conclusions could be made by experimental biochemists analyzing the mutants, our tool can quantify the effect of various amino acids at each position on the activity of the enzyme. The tailored workflow that we created for protein/enzyme engineering is provided with extra modules for the analysis and visualization of a mutant dataset. One of these provides 5-fold cross-validation on the whole dataset (**Supplementary Fig. 19c**). K-fold cross-validation assesses the predictivity of machine learning models on an independent dataset (**Methods**). An average of 0.65 (Pearson $R^2$) for 5-fold cross-validation was achieved on this dataset. The other module plots the performance of the model on a single test set, 20% of the whole data, after being trained on the other 80%. **Supplementary Fig. 19d** shows predicted values of the test set versus their measured values for the mutants dataset which supports an existing correlation between predicted and actual values. Our enzyme engineering notebook also enables continuing with active learning cycles from an imported dataset (**Supplementary Fig. 20**). One can also start from scratch to engineer an enzyme by following a procedure similar to the examples shown in this study.

Preparation of the dataset for enzyme mutants
Data for the iterative saturation mutagenesis of MeOXC was acquired as described previously[7]. Briefly, a three-enzyme cascade was employed to turn over formaldehyde and formyl-CoA to glyoxylate. The last step in the cascade formed hydrogen peroxide, which was used to convert Ampliflu Red to resorufin, allowing fluorometric detection. Mutants were evaluated by two parameters. First, the maximal rate of signal production was determined (parameter A), and second, the final amount of signal produced after 2 hours of runtime was measured (parameter B). The assigned objective function to each mutant is the product of parameters A and B. All values were normalized to the wildtype. If possible, the specific mutations were matched to the dataset. Where no sequencing data was available, the average signal was determined and assigned to the missing mutants.

Supplementary Tables:

**Supplementary Table 1. Summary of METIS optimization and prediction Google Colab packages.** In this study, we provided two modular Google Colab notebooks that can be adjusted for different applications as shown in the study. These two notebooks differ in applications, query strategy, inputs and outputs.

| Notebook | Application | Query Strategy | Input/start | Output |
|---|---|---|---|---|
| METIS optimization | -Optimization of an objective function, i.e., a composition, pathway, genetic construct/circuit | -Maximize the objective function as well as the standard deviation of ensemble regressor | -Existing data<br>OR<br>-Start with randomly generated compositions | -Compositions of categorical and/or quantitative factors that lead to higher yields |
| METIS prediction | -Prediction of an objective function, i.e., a composition, pathway, genetic construct/circuit | -Maximize only the standard deviation of ensemble regressor | -Existing data<br>OR<br>-Start with randomly generated compositions | -A trained model that predicts the objective function of given compositions |

**Supplementary Table 2: The removed conditions after the first 3 rounds of yield active learning.**

| Compounds | Removed values |
|---|---|
| HEPES | 25 |
| MgCl2 | 22.5, 25 |
| Creatine P | 80, 100 |
| Bicarbonate | 100 |
| Formate | 100 |
| B12 | 0.6, 0.8, 1.0 |
| pco | 0.1914, 0.3827, 0.669725, 1.243775 |
| hbs | 1.06785, 2.1357, 3.737475, 6.941025 |
| mco | 1.36885, 2.7377, 4.10655, 5.4754, 6.84425, 9.58195, 13.6885, 19.1639 |
| fdh | 1.456, 2.912, 5.824, 10.192 |
| gor | 0.55235, 1.1047, 1.65705, 2.2094 |

**Supplementary Table 3: Timepoint assays of 7 selected conditions with their active learning yield (AL yield) and efficiency (AL efficiency) values.** Blue, day 4, condition 29, pH 7.8; orange, day 7, condition 15, pH 7.8; red, day 7, condition 76, pH 7.2; black, control, pH 7.6. green, day 8, condition 17, pH 7.2; lavender, day 5, condition 17, pH 7.4; burgundy, day 5, condition 55, pH 7.8.

| | HEPES | MgCl2 | CP | Bicarb. | Form. | CoA | B12 | ATP | NADPH | pco | ccr | epi | mcm | scr | ssr | hbs | hbd | ecm | mco | mch | mcl | kat | fdh | ca | gor | ck | **Manual Yield (3h)** | **AL yield** | **AL Efficiency** |
|---|---|---|---|---|---|---|---|---|---|---|---|---|---|---|---|---|---|---|---|---|---|---|---|---|---|---|---|---|---|
| blue | 75 | 12.5 | 5 | 10.0 | 40 | 0.5 | 0.1 | 10 | 3.75 | 2.3 | 2.2 | 1.5 | 1.5 | 7.0 | 4.4 | 0.5 | 1.5 | 2.9 | 46.5 | 0.3 | 14.7 | 1.6 | 13.1 | 0.0 | 4.4 | 2.7 | 2262.5 | 2869.6 | 26.8 |
| orange | 75 | 12.5 | 60 | 2.5 | 20 | 0.4 | 0.0 | 3 | 3.75 | 3.1 | 1.9 | 0.7 | 2.9 | 3.5 | 1.7 | 0.5 | 0.7 | 1.4 | 26.0 | 0.3 | 3.6 | 3.3 | 30.6 | 0.1 | 5.0 | 0.8 | 2126.8 | 2617.7 | 30.5 |
| red | 150 | 7.5 | 20 | 20.0 | 40 | 0.4 | 0.0 | 9 | 2.50 | 2.3 | 2.2 | 3.7 | 0.3 | 1.7 | 4.4 | 1.6 | 0.4 | 2.6 | 113.6 | 2.3 | 8.4 | 4.9 | 23.3 | 0.1 | 3.9 | 2.7 | 1872.5 | 2586.8 | 14.5 |
| black | 100 | 5.0 | 20 | 50.0 | 20 | 0.5 | 0.1 | 2 | 5.00 | 3.1 | 0.6 | 0.7 | 0.3 | 2.6 | 0.6 | 5.3 | 0.7 | 0.6 | 21.9 | 0.3 | 2.8 | 1.6 | 14.6 | 0.0 | 1.1 | 0.4 | 959.6 | | |
| green | 175 | 10.0 | 10 | 5.0 | 80 | 0.4 | 0.1 | 9 | 2.50 | 2.3 | 0.6 | 3.0 | 0.3 | 5.2 | 1.1 | 1.6 | 0.4 | 2.3 | 26.0 | 1.4 | 14.7 | 1.6 | 40.8 | 0.1 | 2.8 | 2.0 | 920.2 | 919.9 | 8.7 |
| lavender | 50 | 2.5 | 5 | 2.5 | 5 | 0.4 | 0.1 | 7 | 7.50 | 4.0 | 2.8 | 0.7 | 1.2 | 9.6 | 2.2 | 1.6 | 1.1 | 2.6 | 46.5 | 0.6 | 1.2 | 8.2 | 7.3 | 0.1 | 4.7 | 3.1 | 682.7 | 627.9 | 6.4 |
| burgundy | 50 | 17.5 | 10 | 5.0 | 20 | 4.0 | 0.1 | 5 | 5.00 | 3.1 | 1.5 | 3.7 | 0.9 | 13.1 | 4.4 | 12.3 | 1.8 | 2.9 | 34.2 | 2.0 | 1.6 | 4.9 | 23.3 | 0.0 | 3.6 | 0.8 | 296.6 | 354 | 3.1 |

**Supplementary Table 4: Gradient for the separation of CoA esters.**

| **Time [min]** | **A [%]** | **B [%]** |
|---|---|---|
| 0.0 | 100 | 0 |
| 0.5 | 100 | 0 |
| 6.0 | 96 | 4 |
| 10.0 | 77 | 23 |
| 11.0 | 20 | 80 |
| 12.0 | 20 | 80 |
| 12.1 | 100 | 0 |
| 15.0 | 100 | 0 |

**Supplementary Table 5: Multiple reaction monitoring (MRM) transitions for measurement of CoA esters.**

| Compound | Precursor Ion | Product Ion | Dwell | Fragmentor | Collision Energy | Cell Accelerator Volt. | Polarity |
|---|---|---|---|---|---|---|---|
| Ethylmalonyl-CoA (Quantifier) | 882.1 | 331.2 | 25 | 380 | 41 | 5 | Positive |
| Ethylmalonyl-CoA (Qualifier) | 882.1 | 428 | 25 | 380 | 29 | 5 | Positive |
| Methylsuccinyl-CoA (Quantifier) | 882 | 375.1 | 25 | 380 | 33 | 5 | Positive |
| Methylsuccinyl-CoA (Qualifier) | 882 | 428 | 25 | 380 | 29 | 5 | Positive |
| Mesaconyl-CoA (Quantifier) | 880.1 | 375.1 | 25 | 380 | 25 | 5 | Positive |
| Mesaconyl-CoA (Qualifier) | 880.1 | 428 | 25 | 380 | 35 | 5 | Positive |
| Succinyl-CoA (Quantifier) | 868.1 | 361.1 | 25 | 380 | 35 | 5 | Positive |
| Succinyl-CoA (Qualifier) | 868.1 | 428.1 | 25 | 380 | 35 | 5 | Positive |
| Methylmalonyl-CoA (Quantifier) | 868.1 | 317.1 | 25 | 380 | 41 | 5 | Positive |
| Methylmalonyl-CoA (Qualifier) | 868.1 | 428 | 25 | 380 | 31 | 5 | Positive |
| 4-hydroxybutyryl-CoA (Quantifier) | 854.1 | 347.1 | 25 | 380 | 37 | 5 | Positive |
| 4-hydroxybutyryl-CoA (Qualifier) | 854.1 | 428 | 25 | 380 | 30 | 5 | Positive |
| Crotonyl-CoA (Quantifier) | 836.1 | 329 | 25 | 380 | 33 | 5 | Positive |
| Crotonyl-CoA (Qualifier) | 836.1 | 428 | 25 | 380 | 26 | 5 | Positive |
| Propionyl-CoA (Quantifier) | 824.1 | 317.1 | 25 | 380 | 31 | 5 | Positive |
| Propionyl-CoA (Qualifier) | 824.1 | 428 | 25 | 380 | 28 | 5 | Positive |
| B-methylmalyl-CoA (Quantifier) | 898.1 | 391.1 | 25 | 380 | 39 | 5 | Positive |
| B-methylmalyl-CoA (Qualifier) | 898.1 | 428.1 | 25 | 380 | 33 | 5 | Positive |

**Supplementary Table 6: Gradient for the separation of glycolate.**

| Time [min] | A [%] | B [%] |
|---|---|---|
| 0 | 100 | 0 |
| 4 | 100 | 0 |
| 6 | 0 | 100 |
| 7 | 0 | 100 |
| 7.1 | 100 | 0 |
| 12 | 100 | 0 |

**Supplementary Table 7: Multiple reaction monitoring (MRM) transitions for measurement of glycolate.**

| Compound | Precursor Ion | Product Ion | Dwell | Fragmentor | Collision Energy | Cell Accelerator Volt. | Polarity |
|---|---|---|---|---|---|---|---|
| $^{12}$C-Glycolate (Quantifier) | 75 | 47 | 150 | 380 | 9 | 5 | Negative |
| $^{12}$C-Glycolate (Qualifier) | 75 | 75 | 150 | 380 | 0 | 5 | Negative |
| $^{13}$C-Glycolate (Quantifier) | 77 | 48 | 150 | 380 | 9 | 5 | Negative |
| $^{13}$C-Glycolate (Qualifier) | 77 | 77 | 150 | 380 | 0 | 5 | Negative |

Supplementary Figures:

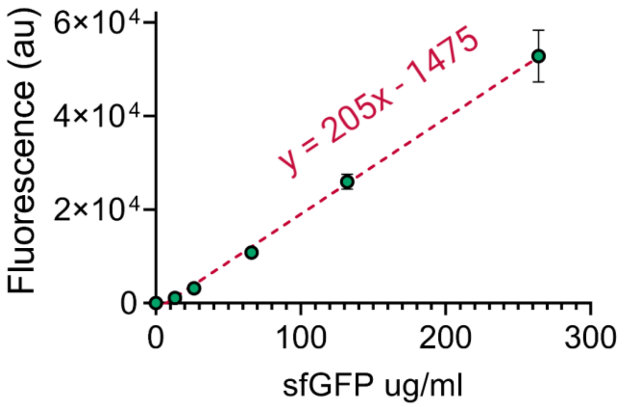


**Supplementary Fig. 1: Standard curve for fluorescence readout of purified sfGfp (26806 Da) at the condition of the reference cell-free reaction.** The experiment was performed in the same plate reader (Tecan Infinite 200 PRO, excitation/emission wavelengths of 485/528 nM and gain = 80). The average of measured Gfp fluorescence for the reference composition[8] is 3849.13 which corresponds to a concentration of 25.97 ug/ml (0.97 µM). This is the amount of sfGfp for the reference that we used to normalize all the data points in **Fig. 1e**. Quick math shows that the points with a yield of ~15 in **Fig. 1e** have a GPF production of ~0.97 $\times$ 15 = 14.5 µM. Compared to a previous optimization and a commercial kit (See the supplementary information of Borkowski *et al.* Figure 8b[2]) we achieved more than 30-fold higher sfGfp production. It should be noted that we used autolysate preparation protocol whereas in the other work, a sonication protocol was used. Moreover, it is important to note that the concentration of plasmid DNA used in this study is 20 nM compared to the 10 nM in the compared work, however, they used a stronger RBS, B0034. The strength of RBS (B0032) in our plasmid is 30% of B0034[9].

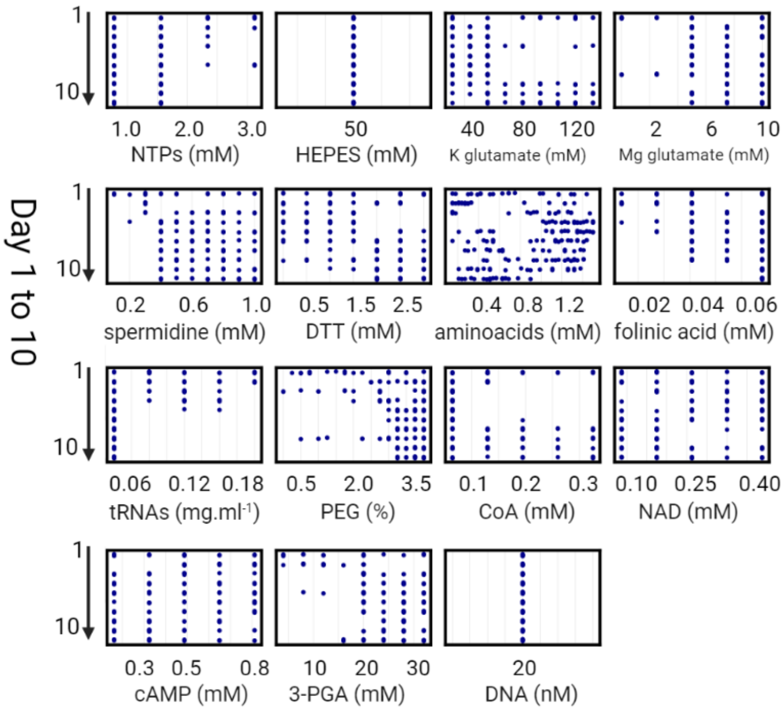


**Supplementary Fig. 2: Concentration variations of each of the *E. coli* cell-free system factors from rounds 1 to 10 of active learning.** These plots show how the concentration of each factor varied from day 1 to 10 through suggestions of the model aiming to increase the yield. The features' importance (**Fig. 1f**), and yield dependencies (**Fig. 1g**) could be analyzed altogether with these plots.

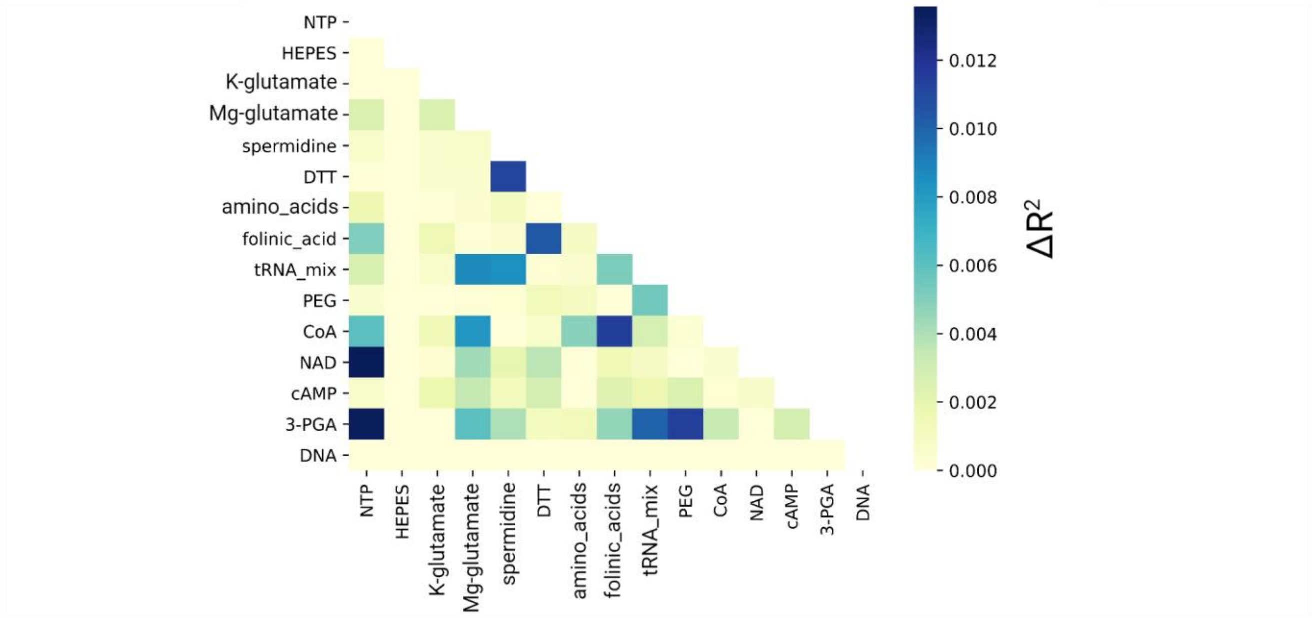


**Supplementary Fig. 3: Mutual interactions between every two factors for active learning of the *E. coli* cell-free system.** This plot shows the calculated mutual interactions (**Methods**) between every two factors, a useful analysis of the active learning dataset helping to study the system. This module is also integrated into the modular workflow.

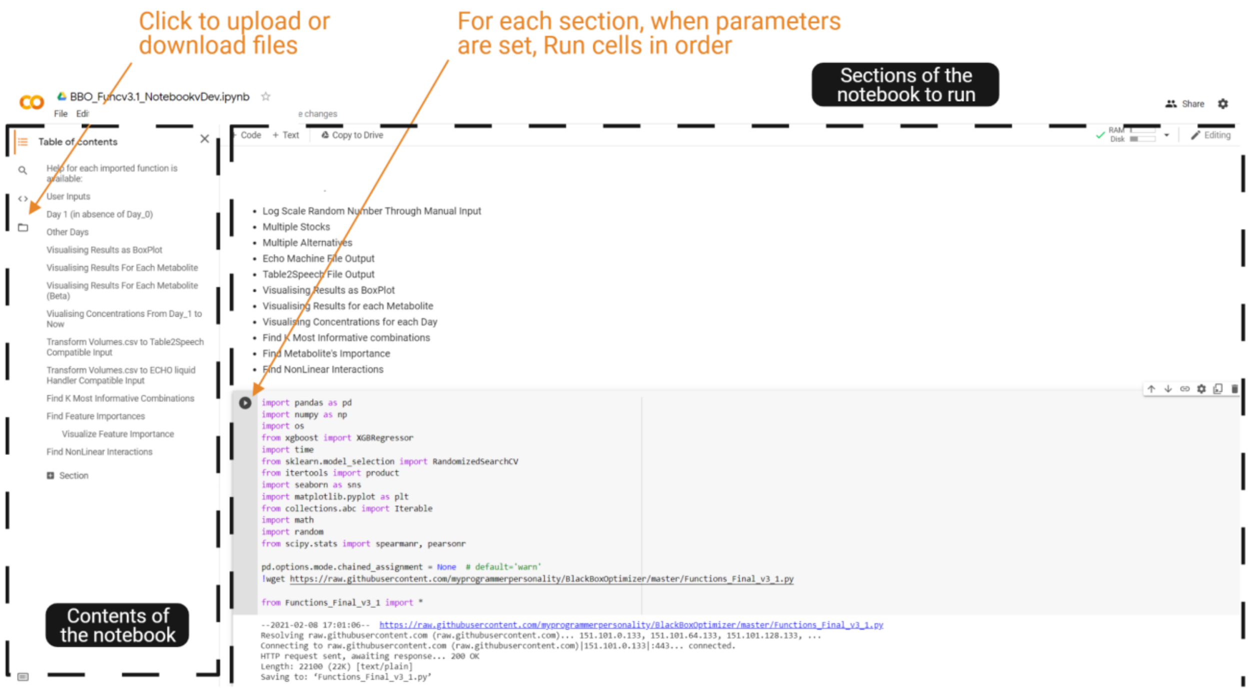


**Supplementary Fig. 4: A screenshot of the Google Colab notebook for interacting with the workflow.** At the left side of the page, the folder icon (shown by the arrow) is where files should be uploaded (Results) or downloaded (those that the user generated during the usage such as Volume, Concentration, Features' importance, K most informative, as well as figures as .png or .svg). When the run time is over, the user should reupload the files. The main body of the tool is where different cells (modules) of the code are accessible that should be run in order by clicking on the run icon (shown by the arrow). Except for the first round for which there is no results or input file, the files should be uploaded before starting to run the cells. For more details see **Code availability** to open (after making a personal copy) any of provided notebooks in your browser.

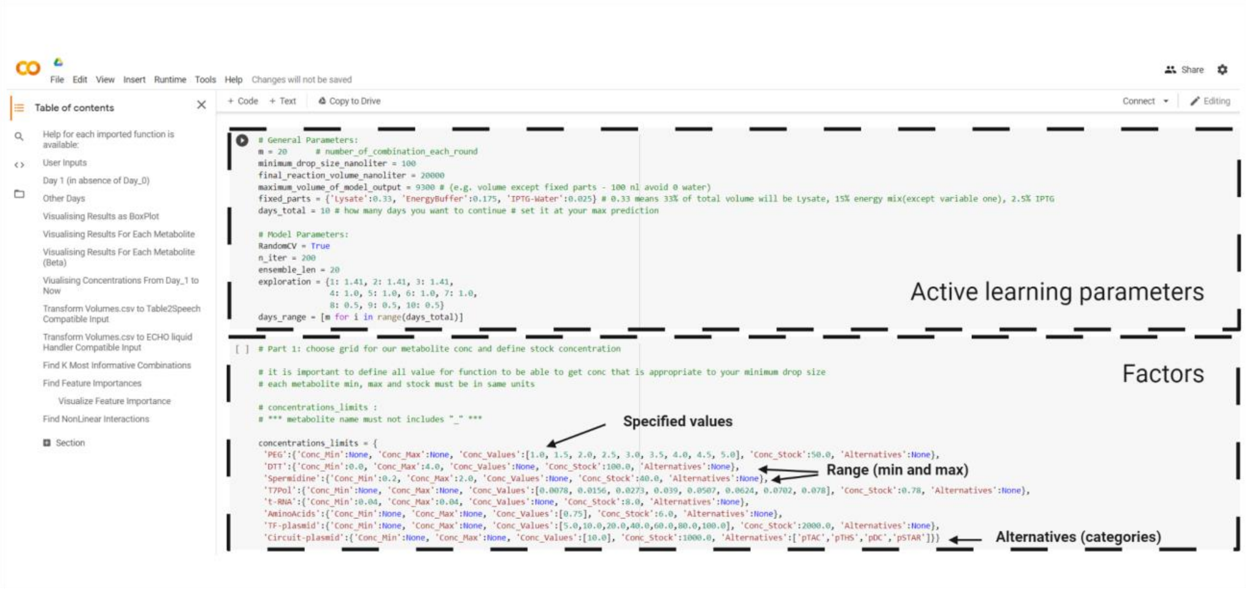


**Supplementary Fig. 5: A screenshot of the sections of active learning parameters and factors.** These two sections appear at the top of the notebook. These cells come after the sections for python classes/modules that should run beforehand. See **Supplementary Note 2** for detailed explanations.

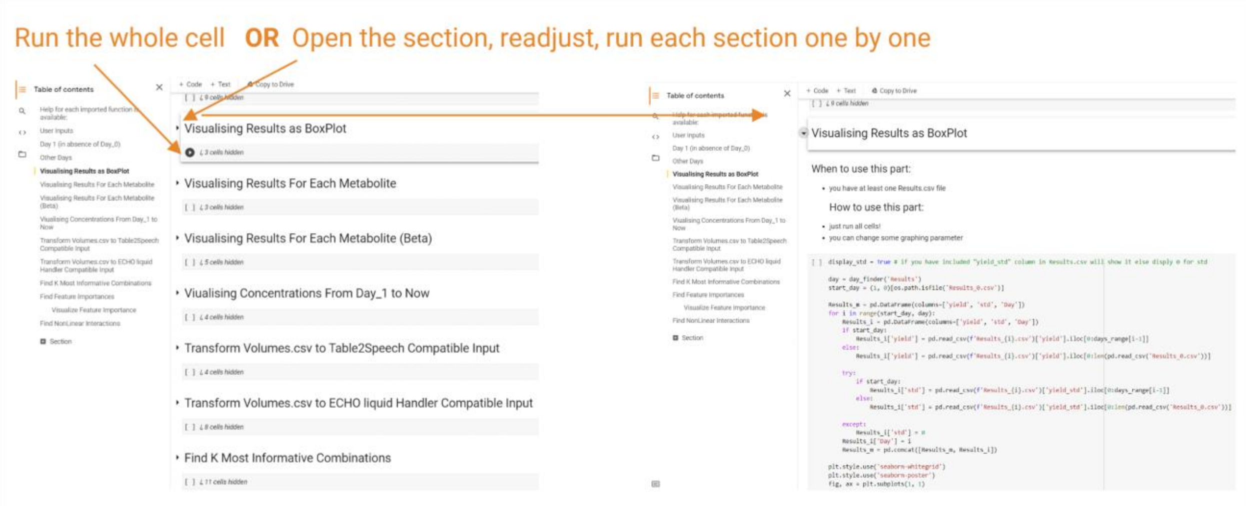


**Supplementary Fig. 6: A screenshot of different modules of the workflow.** The modules that a user aims to use can be run all at once or the user can open individual code sections/tabs and make changes for example in the plot size, color, etc.

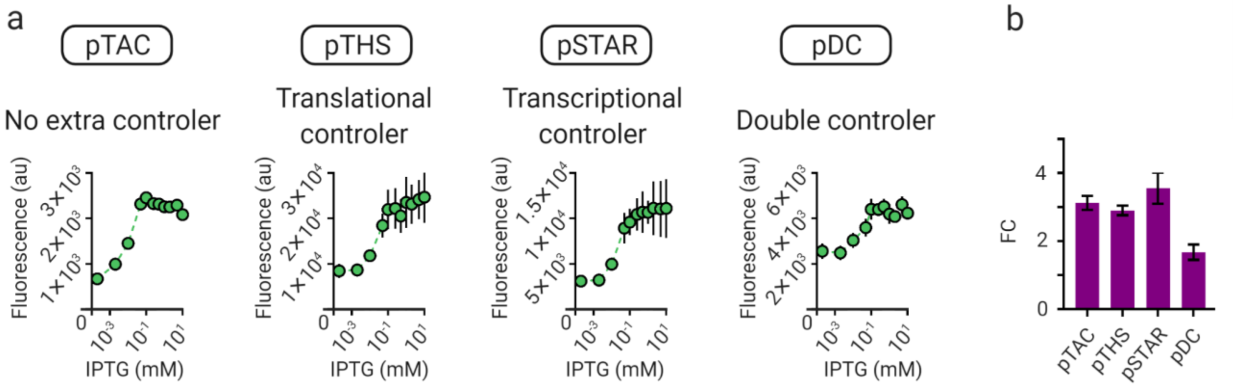


**Supplementary Fig. 7: The behavior of *LacI* gene circuits in the cell-free protein expression system.** These results presented in Greco *et al.*[10] as the "rate" in the protein production; however, here, we plotted the "amount" of protein produced after 6 h of incubation at 30 °C. **(a)** IPTG dose-response curve for SLC and MLC constructs. **(b)** The fold-change (FC) value of the plots in **(a)** between 0 and 10 mM of IPTG.

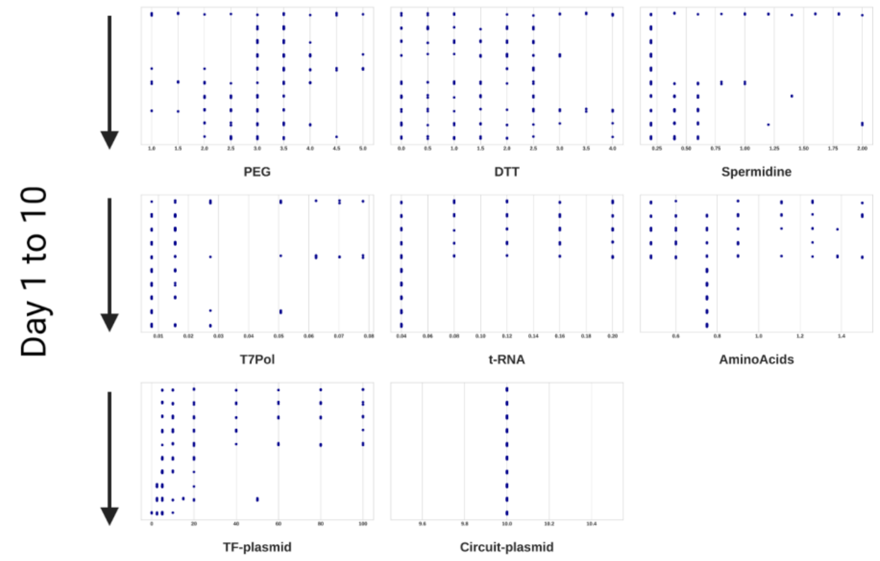


**Supplementary Fig. 8: Variation of the concentration of factors in 10 rounds of active learning for *LacI* gene circuits optimization.** These plots show suggestions of different concentrations of each factor by the model aiming to improve the objective function.

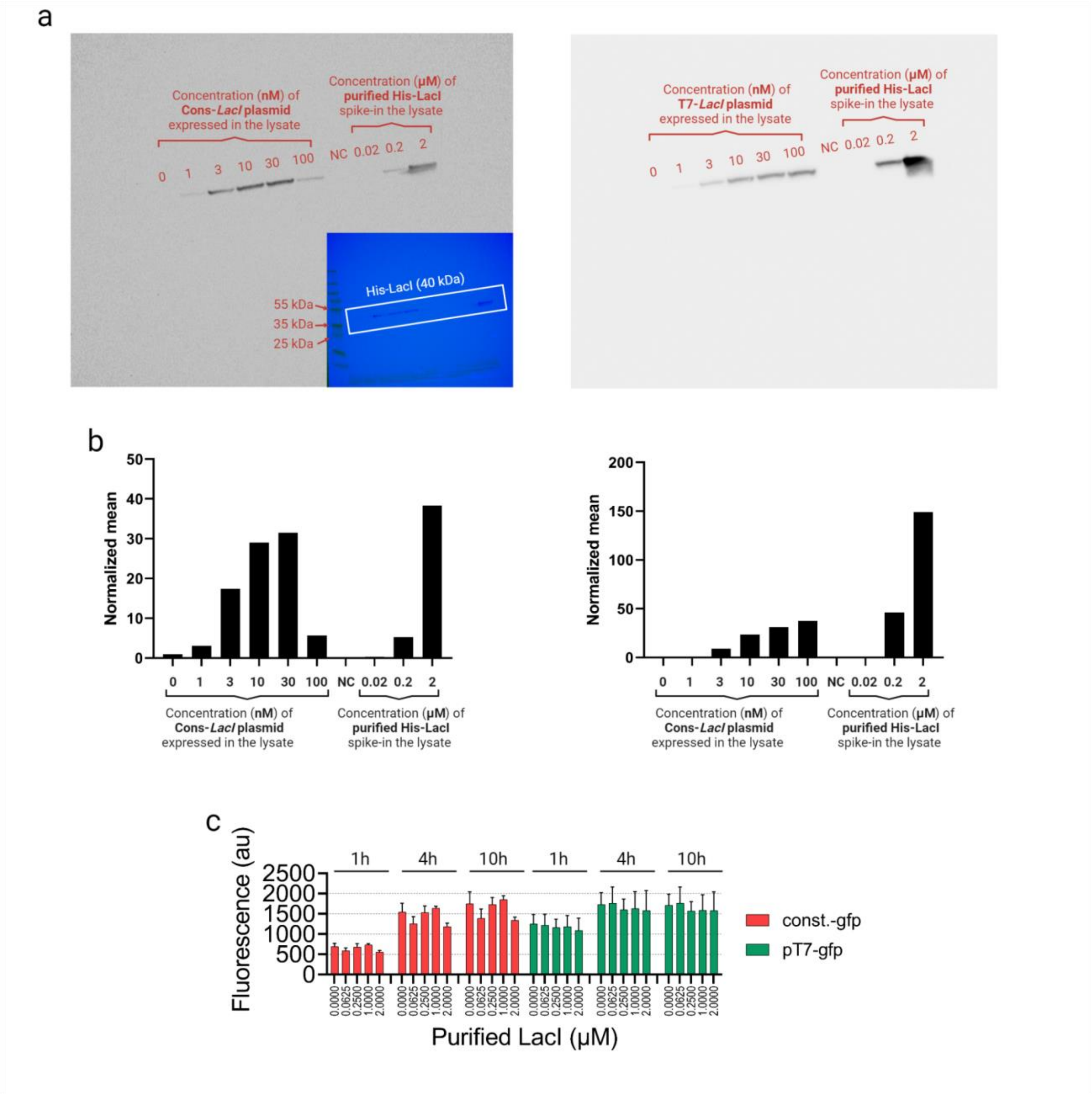


**Supplementary Fig. 9: Gfp fluorescence level is affected by expression competition with *LacI*, not by interference by the LacI protein itself. a)** Western chemiluminescence detection of LacI protein level produced in cell-free reactions starting with different concentrations of the *LacI* plasmid. The inset shows an overlay of the chemiluminescent signal and a bright image of the membrane, where the ladder is visible. **b)** Quantification of the mean intensity of LacI bands in **(a)** using imageJ software. These data show that all cell-free reactions produce less than 2 µM of the LacI protein. **c)** Gfp fluorescence from cell-free reactions with different concentrations of purified LacI-6xHis protein included indicates that the LacI protein itself does not affect *Gfp* expression level at 2 µM and below.

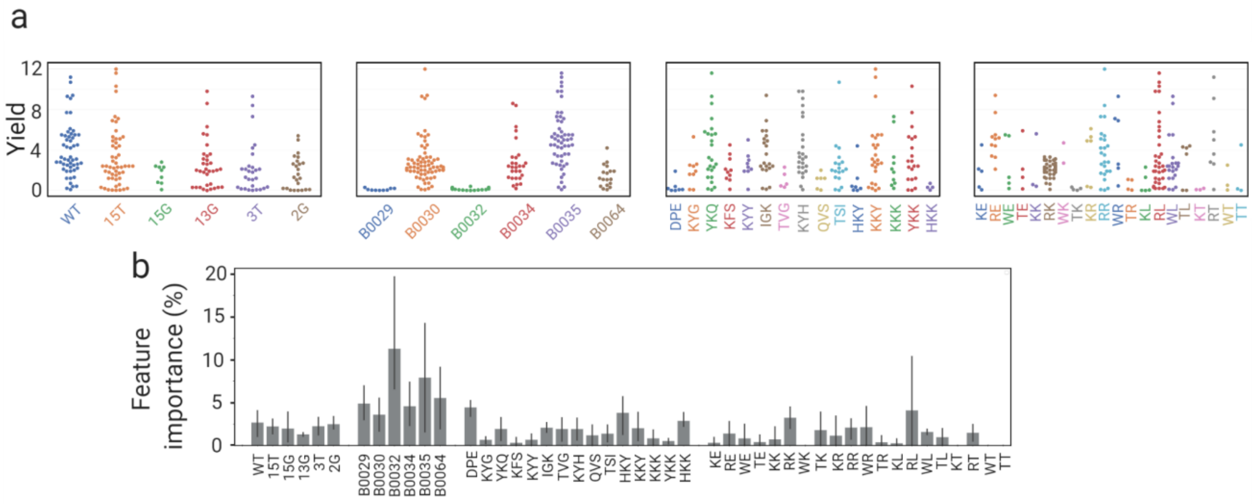


**Supplementary Fig. 10: Analysis of the active learning data for the transcription and translation unit. (a)** Distribution of alternative factors within the yield of 200 combinations. **(b)** The feature importance percentage of each condition for the model to assign predicted yield values. The bars and error bars are the average and standard deviation of the importances in 4 days

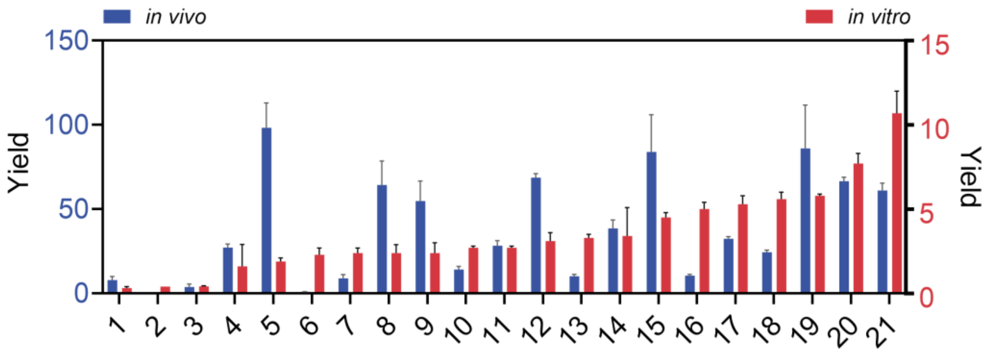


**Supplementary Fig. 11: Cell-free versus *in vivo* yields of the 20 most informative combinations for the transcription & translation unit.** The bar plot representation of results (average and standard deviation of triplicates) shown in **Fig. 4e**.

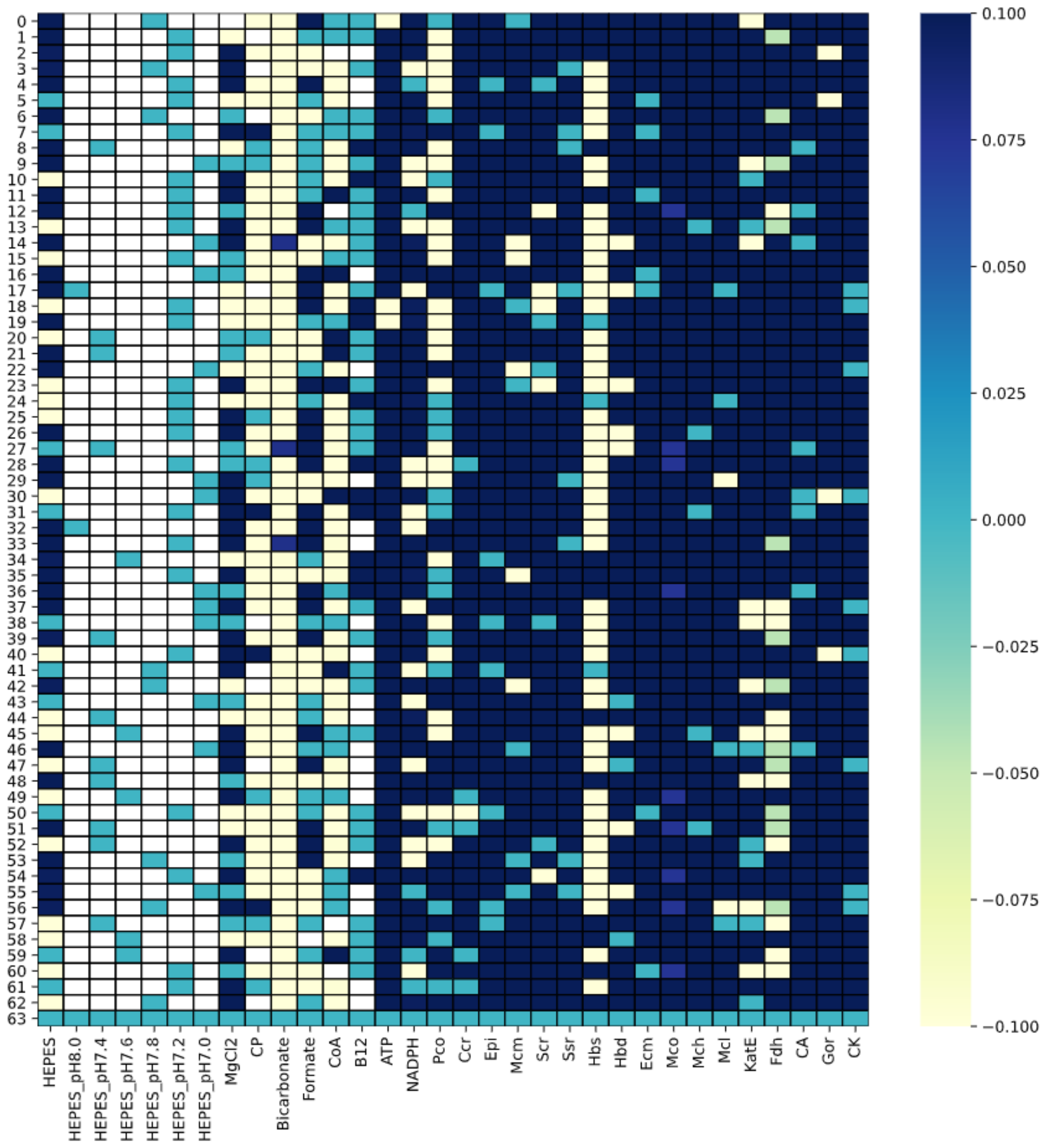


**Supplementary Fig. 12: Scaled factors heatmap of top 10% yields in CETCH yield (glycolate, round 1-5) active learning.** The concentration range for factors was scaled to the same range.

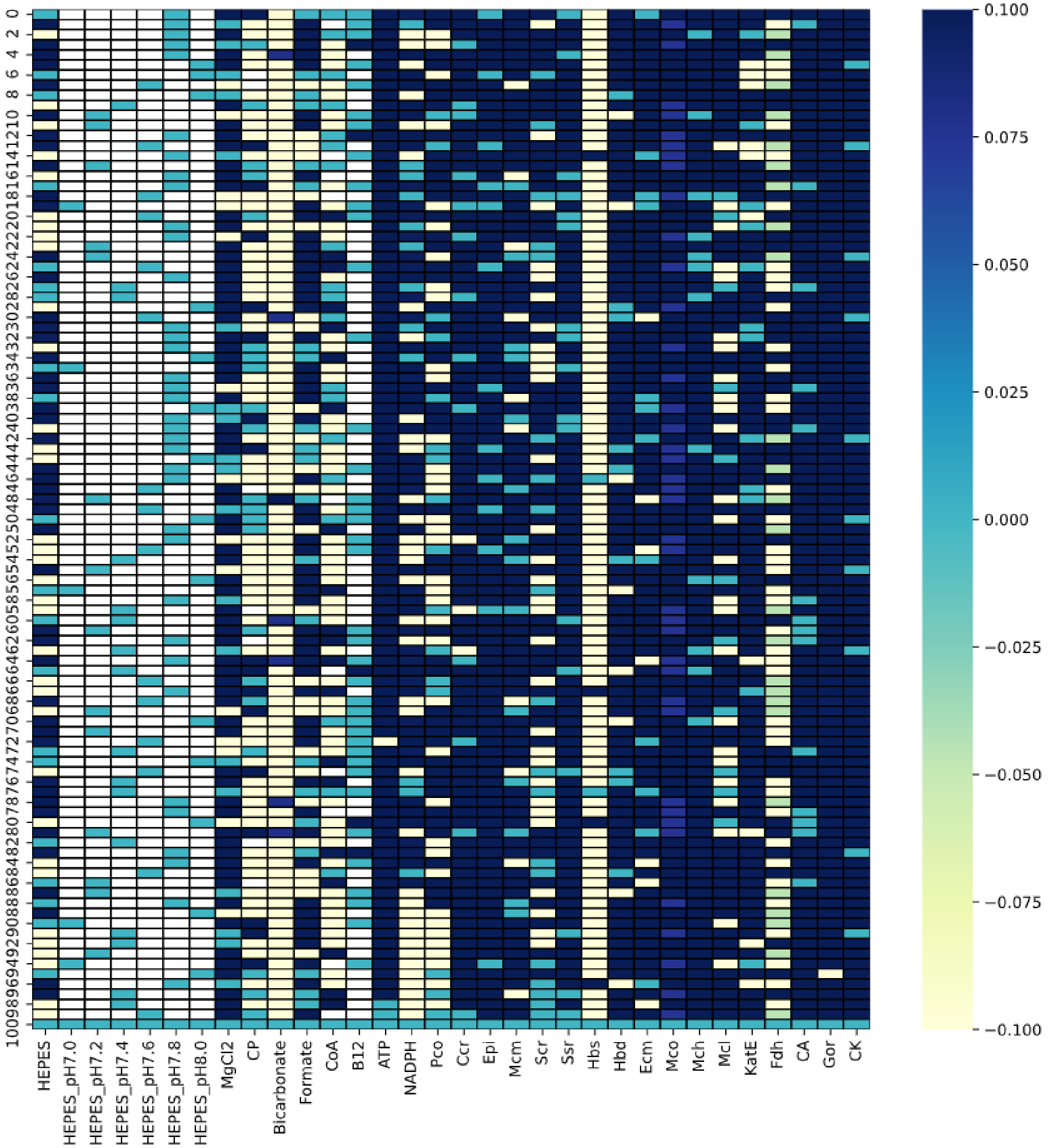


**Supplementary Fig. 13: Scaled factors heatmap of top 10% yields in CETCH efficiency (glycolate concentration divided by the total enzyme concentration, round 1-8) active learning.** The concentration range for factors was scaled to the same range.

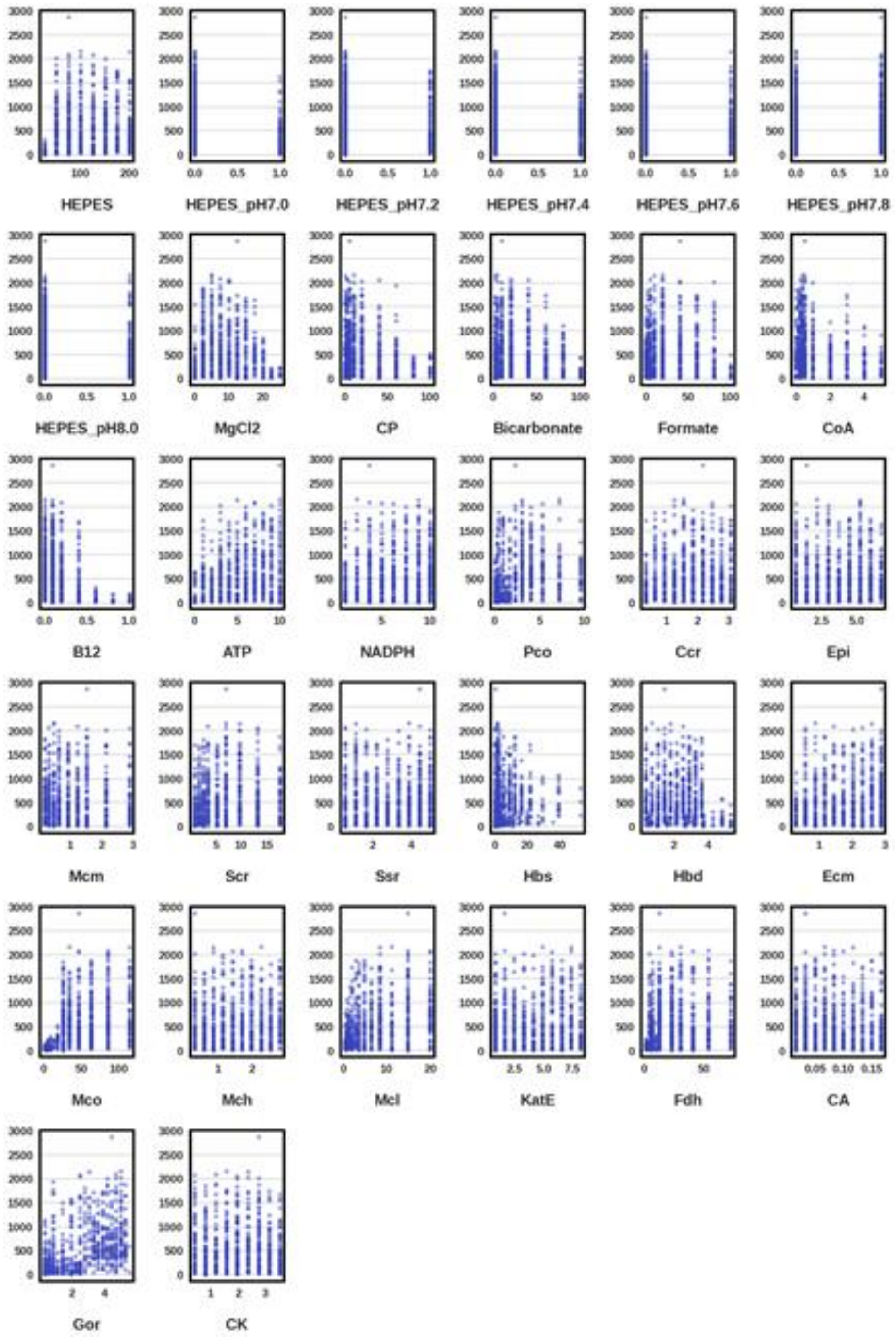


**Supplementary Fig. 14: Distribution of measured yield (glycolate, round 1-5) values within the ranges of each factor.** Distribution of all factors within the yield of 5 rounds.

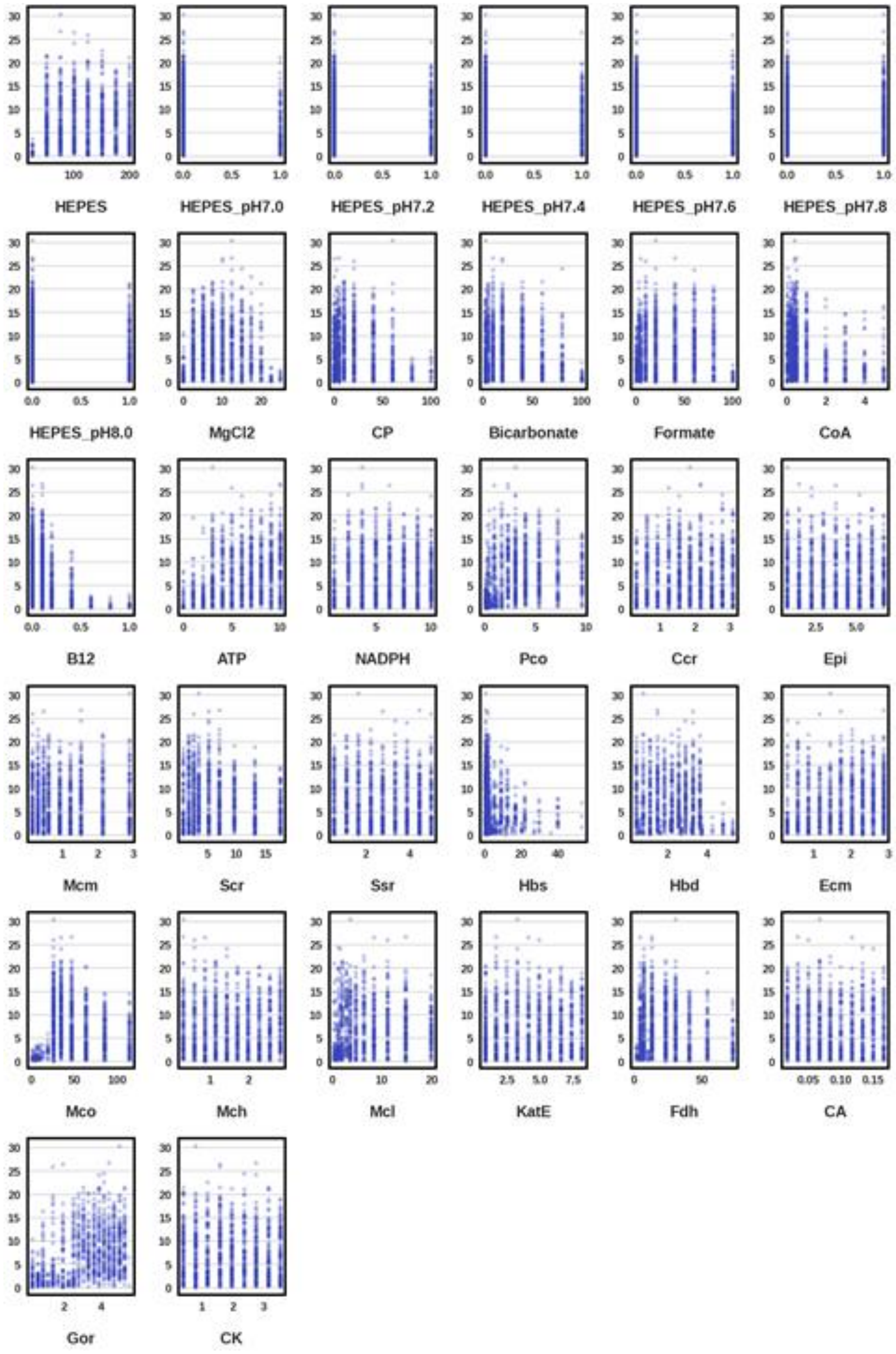


**Supplementary Fig 15: Distribution of measured efficiency (glycolate concentration divided by the total enzyme concentration, round 1-8) values within the ranges of each factor.** Distribution of all factors within the efficiency of 8 rounds.

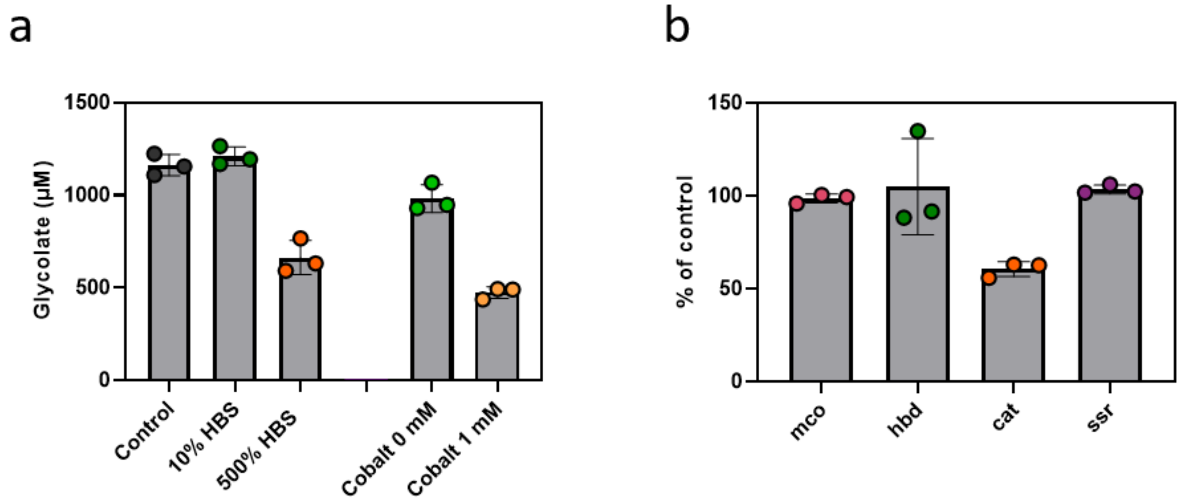


**Supplementary Fig. 16: Assays for testing newly purified enzyme batches after round 2 (b) and for testing different concentrations of hbs and cobalt (a). (a)** The assays were done as described in material and methods (Assays for different concentrations of hbs and cobalt). The shown glycolate concentrations are from the sample taken after 180 min. The control contained 0.1 mM Coenzyme $B_{12}$, whereas the assays with 0 mM and 1 mM cobalt did not contain any Coenzyme $B_{12}$. **(b)** Test of old batches of mco, hbd, cat and ssr. After recognizing an increase in the product yield of our control after round two, we tested the old stocks of the enzymes which were purified freshly for round three in combination with the new enzymes. We could identify the old batch of catalase as the reason for the lower yield. The values of the grey bars are the means of three replicates (dots) and the error bars represent the standard deviations.

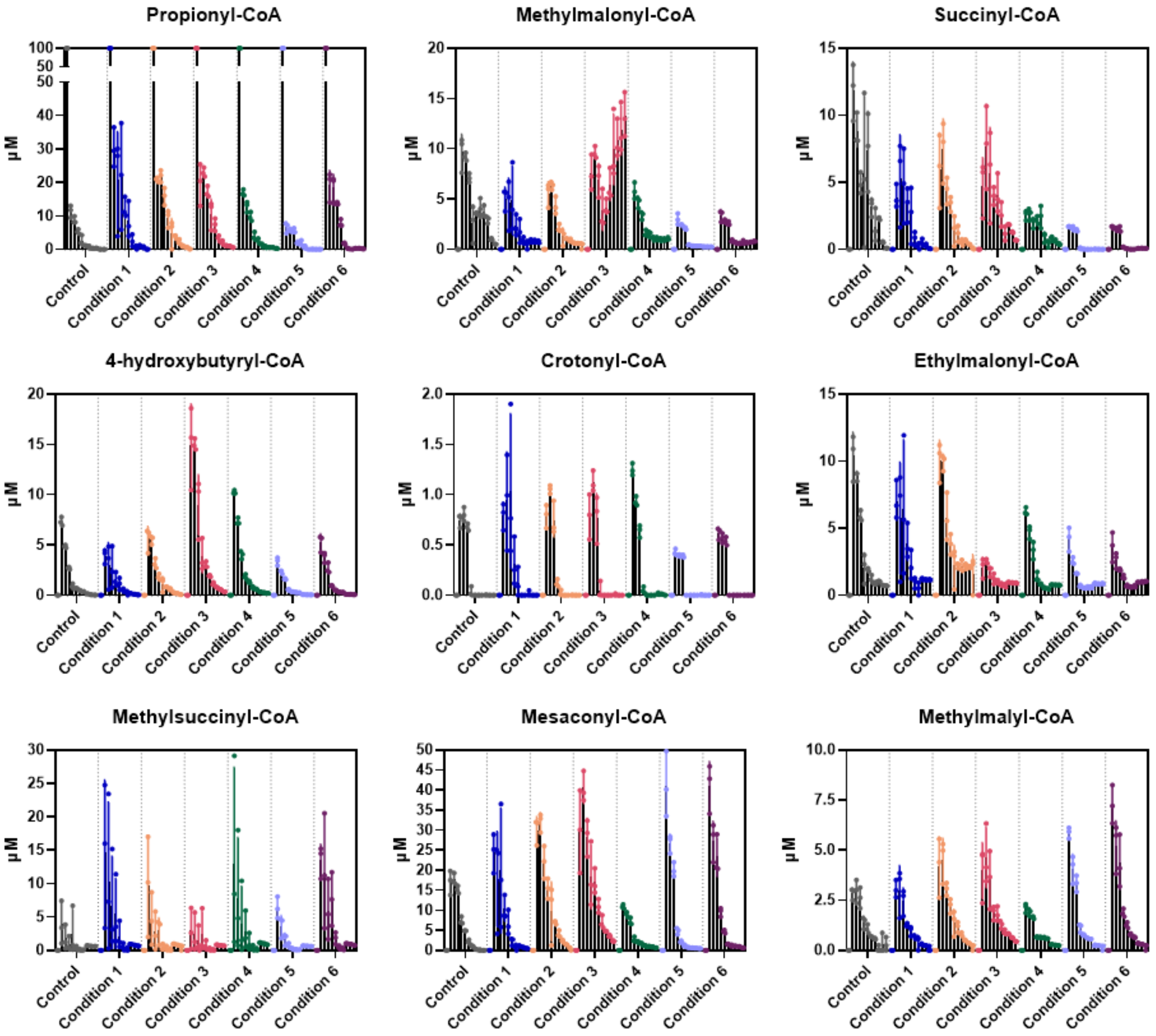


**Supplementary Fig. 17:** Quantified CoA-esters from selected assay conditions. The values for time point 0 are not measured, the reactions were started with 100 µM propionyl-CoA. The values of the black bars are the means of three replicates (dots) and the error bars represent the standard deviations. For details see <u>LC-MS analysis of CoA esters.</u>

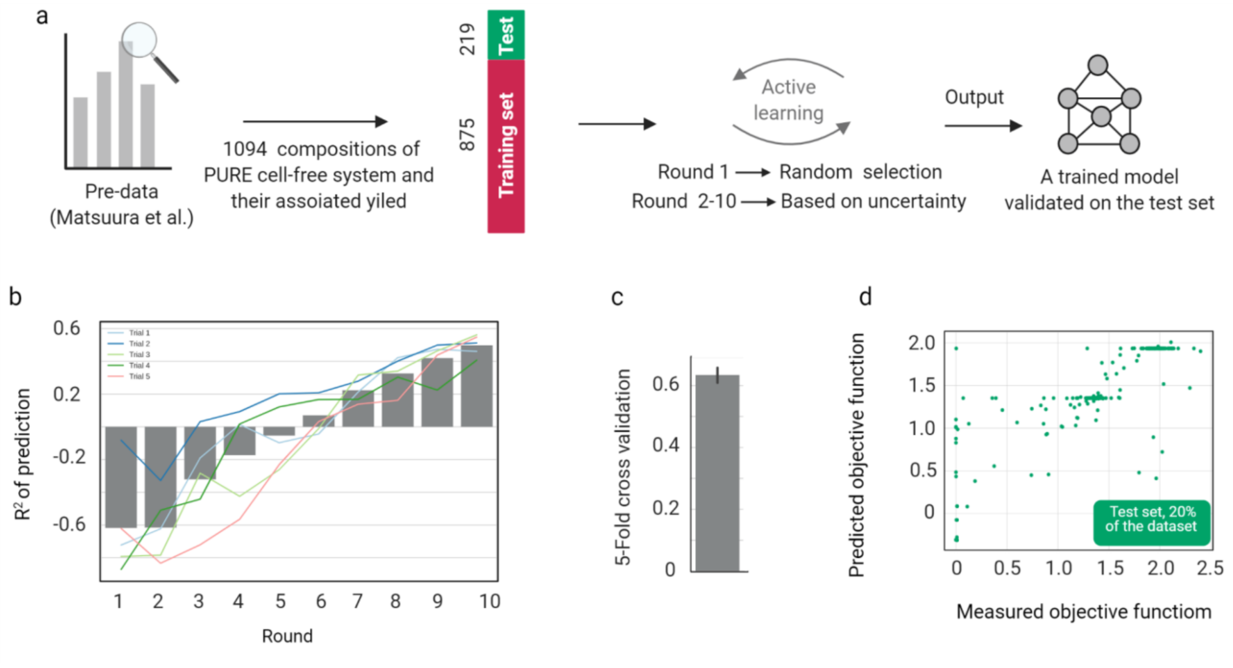


**Supplementary Fig. 18: Application of METIS for the prediction of a PURE cell-free system dataset. (a)** We took this dataset from a recent study in which they varied recombinant proteins and buffer compositions, adapted it as a Results.csv file. The dataset was divided into two subsets 80% for training and 20% for testing. (b) During 10 rounds of active learning, the model was assessed by the initial test set and the $R^2$ of the prediction improved over rounds. See **Supplementary Note 6** for a detailed explanation of the process. **(c)** 5-fold cross-validation to evaluate the average performance of the model on the whole dataset. **(d)** a single round of validation of a model on the test set.

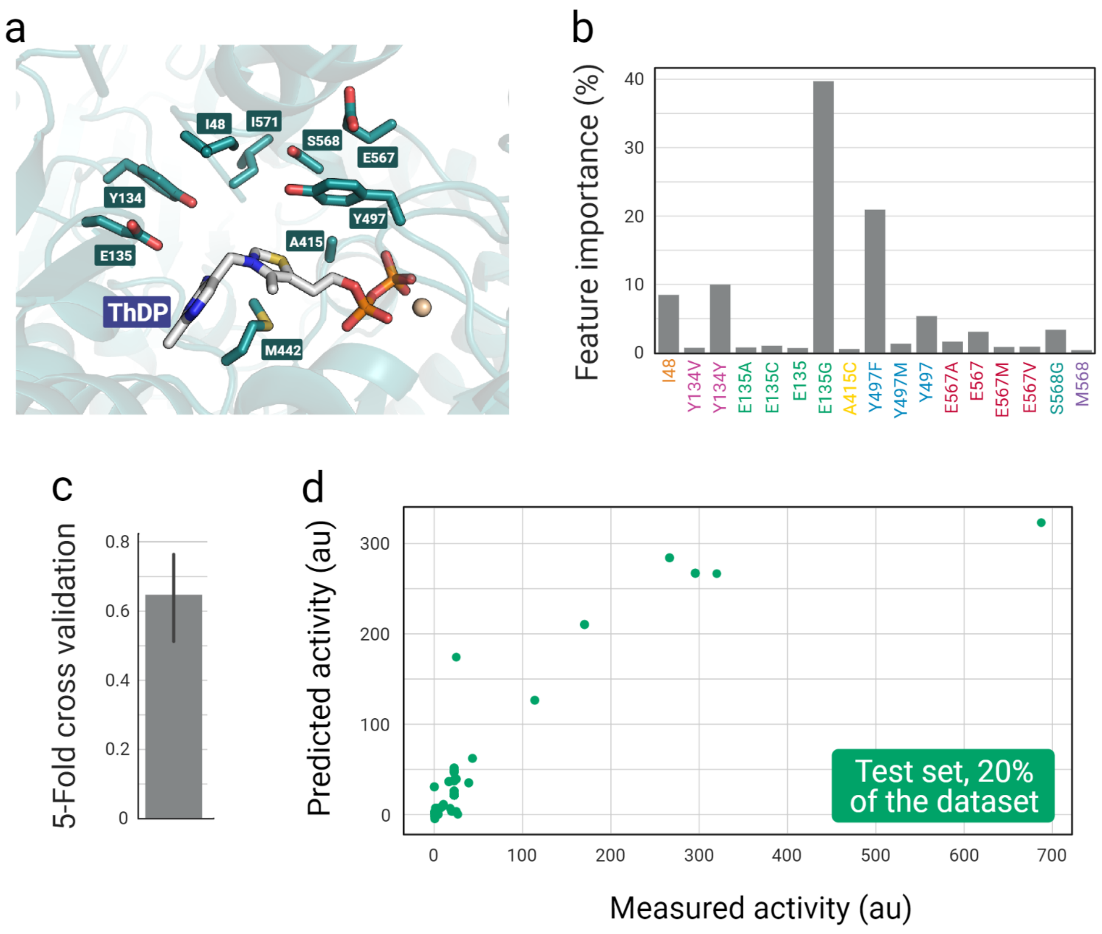


**Supplementary Fig. 19: Simulations on a dataset of 847 mutants of oxalyl-CoA decarboxylase enzyme to showcase the application of METIS workflow for combinatorial enzyme engineering. (a)** The active site of oxalyl-CoA decarboxylase with surrounding residues annotated and the covalently bound cofactor thiamine diphosphate (ThDP). **(b)** Calculated features' importance by the model on the imported dataset that define how big the effect of each condition is if the model suggests new amino acid combinations. **(c)** 5-fold cross-validation, a common approach to assess the prediction power of machine learning models. **(d)** Predicted values on a single unseen test set (20% of the whole dataset) versus their measured values. (see also **Supplementary Note 7**)

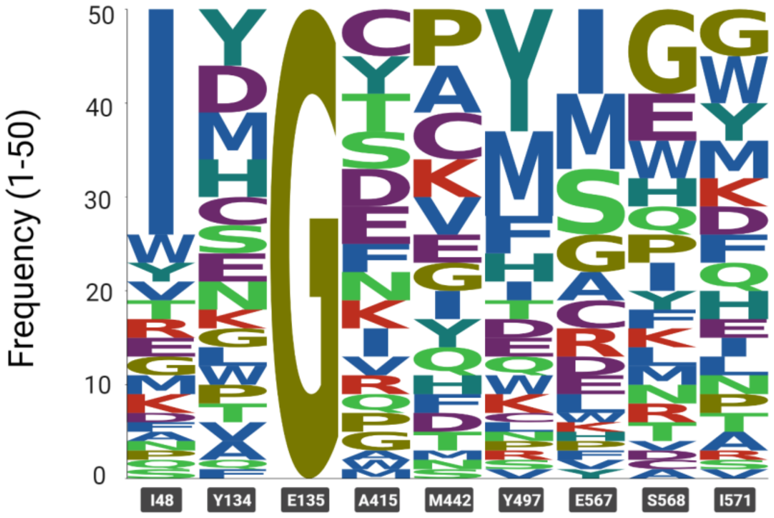


**Supplementary Fig. 20: Distribution of amino acids in each position of the oxalyl-CoA decarboxylase active site predicted by the model out of 50 suggestions.** This shows an example in case a user aims to synthesize or clone new enzyme sequences and test a round of combinatorial mutants.
